# Mechanisms of dual modulatory effects of spermine on the mitochondrial calcium uniporter complex

**DOI:** 10.1101/2023.06.06.543936

**Authors:** Yung-Chi Tu, Fan-Yi Chao, Ming-Feng Tsai

## Abstract

The mitochondrial Ca^2+^ uniporter mediates the crucial cellular process of mitochondrial Ca^2+^ uptake, which regulates cell bioenergetics, intracellular Ca^2+^ signaling, and cell death initiation. The uniporter contains the pore-forming MCU subunit, an EMRE protein that binds to MCU, and the regulatory MICU1 subunit, which can dimerize with MICU1 or MICU2 and under resting cellular [Ca^2+^] occludes the MCU pore. It has been known for decades that spermine, which is ubiquitously present in animal cells, can enhance mitochondrial Ca^2+^ uptake, but the underlying mechanisms remain unclear. Here, we show that spermine exerts dual modulatory effects on the uniporter. In physiological concentrations of spermine, it enhances uniporter activity by breaking the physical interactions between MCU and the MICU1-containing dimers to allow the uniporter to constitutively take up Ca^2+^ even in low [Ca^2+^] conditions. This potentiation effect does not require MICU2 or the EF-hand motifs in MICU1. When [spermine] rises to millimolar levels, it inhibits the uniporter by targeting the pore region in a MICU-independent manner. The MICU1-dependent spermine potentiation mechanism proposed here, along with our previous finding that cardiac mitochondria have very low MICU1, can explain the puzzling observation in the literature that mitochondria in the heart show no response to spermine.

## Introduction

The mitochondrial calcium uniporter (hereafter referred to as the uniporter) is a multi-subunit Ca^2+^ channel that mediates mitochondrial Ca^2+^ uptake. When intracellular Ca^2+^ signals arrive at mitochondria, the uniporter becomes activated to rapidly transport Ca^2+^ into the mitochondrial matrix to modulate the magnitude and frequency of these Ca^2+^ signals^1, 2^. The imported Ca^2+^ can also raise matrix [Ca^2+^] to stimulate multiple Ca^2+^-dependent dehydrogenases in the TCA cycle and enhance oxidative phosphorylation^1, 2^. However, entry of too much Ca^2+^ can induce the mitochondria permeability transition, leading to apoptotic cell death^1, 2^. Malfunction of the uniporter has been implicated in a wide range of pathologies^3, 4^, including cardiac ischemia-reperfusion injury, heart failure, neurodegeneration, and cancer metastasis, among others. Considering the uniporter’s critical importance in pathophysiology, it is important to understand how its activity is regulated.

It has been known for decades that the uniporter is regulated by cytoplasmic Ca^2+^. Following the discovery of uniporter genes in the early 2010s^5-8^, extensive studies have established an “occlusion” mechanisms underlying such Ca^2+^ regulation^9-20^. As summarized in Fig. 1A, the uniporter is inactive in resting cellular [Ca^2+^] (∼100 nM). This is because a MICU1 subunit, which can form a homodimer or heterodimerize with MICU2 in the intermembrane space (IMS), blocks the IMS entrance of the uniporter’s Ca^2+^ pore formed by the MCU subunit. When cytoplasmic Ca^2+^ signals elevate IMS [Ca^2+^], Ca^2+^ binding to MICU1 would cause MICU1 separation from MCU to open the pore, thus leading to Ca^2+^ activation of the uniporter. In this Ca^2+^-activated state, MICU1 remains bound to an EMRE subunit via electrostatic interactions. This interaction ensures unfailing MICU1 association within the uniporter complex, so that once the Ca^2+^ signal is over, MICU1 can rapidly block the Ca^2+^ pore to close the uniporter.

**Figure 1.**
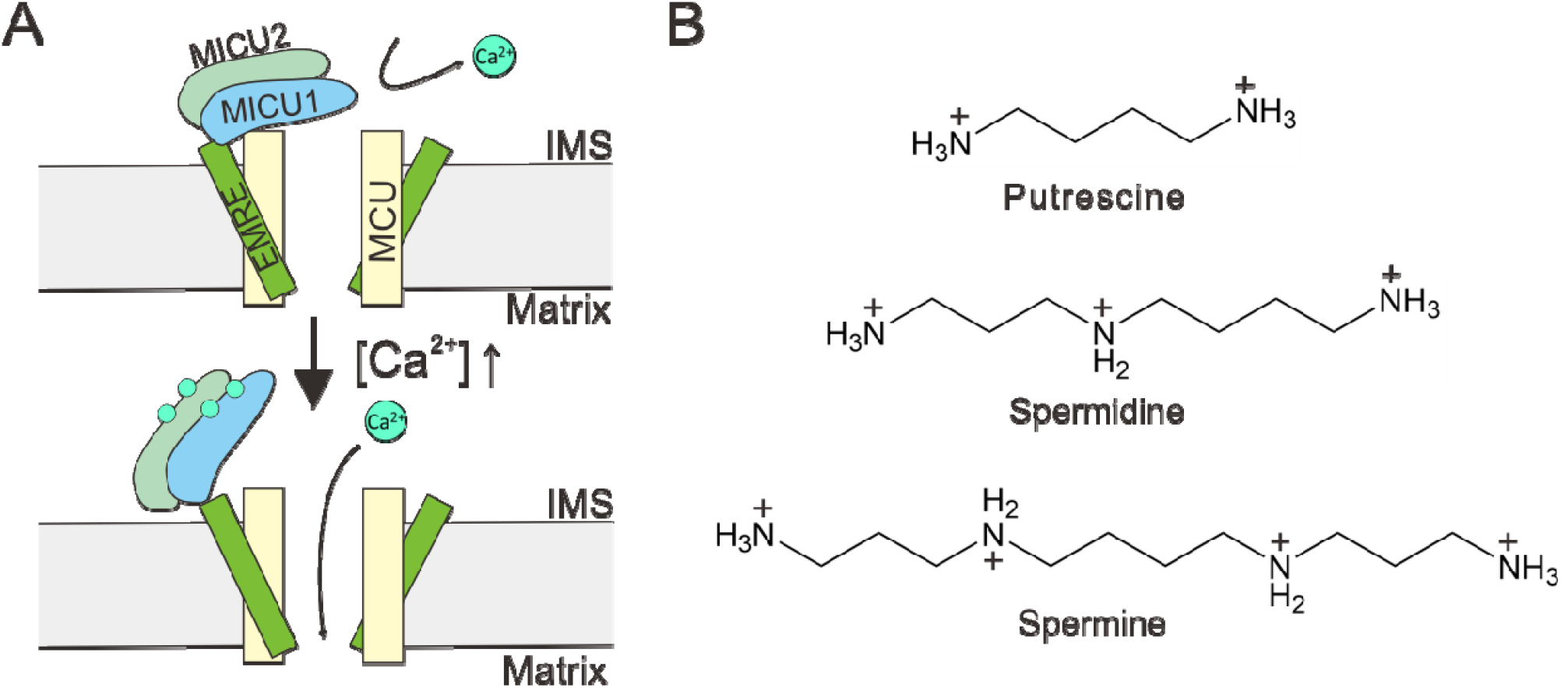
Regulation of the mitochondrial calcium uniporter. **(A)** A molecular model underlying Ca^2+^-dependent uniporter activation. **(B)** Chemical structures of common biological polyamines.

Here, we investigate a possible uniporter regulator, spermine, which along with spermidine and putrescine (Fig. 1B) are important biological polyamines ubiquitously present in animal cells, playing critical roles in cell growth, differentiation, protein synthesis, and apoptosis^21-23^. Spermine, which is a positively charged molecule, is known to modulate the activity of many cation channels by blocking the channel pore. For example, it confers the inward-rectification property of multiple inward-rectifier K^+^ (K_ir_) channels and AMPA-/kainate-type glutamate-receptor channels by entering into the pore from the intracellular side to block outward currents upon membrane depolarization^24-29^. Block of cyclic nucleotide-gated channels^30^, voltage-gated Na^+^ channels^31, 32^, and transient receptor potential channels^33, 34^ has also been reported.

Interestingly, Nicchitta and Williamson^35^ reported in 1984 that spermine can enhance the ability of mitochondria to buffer extramitochondrial Ca^2+^. In particular, they showed that isolated liver mitochondria, or mitochondria in permeabilized hepatocytes, can only reduce extramitochondrial [Ca^2+^] ([Ca^2+^]_ex_) to a steady-state level of 0.5–1 µM. However, adding spermine causes mitochondria to absorb more Ca^2+^, thus lowering [Ca^2+^]_ex_ to 0.2–0.5 µM. It was also shown that spermidine can similarly improve mitochondrial Ca^2+^ buffering, albeit with ∼5-fold lower efficacy and potency, but putrescine is ineffective. These initial observations have since been verified in multiple laboratories^36-42^.

It has been speculated that spermine enhances mitochondrial Ca^2+^ buffering by stimulating mitochondrial Ca^2+^ uptake, which is mediated by the uniporter. However, kinetic analyses have led to puzzling results. Some investigators found that spermine increases the rate of mitochondrial Ca^2+^ uptake^39^, but others observed only inhibitory effects^36, 37, 43^. Moreover, some groups reported more complicated phenomena, with spermine increasing mitochondrial Ca^2+^ uptake rate when [Ca^2+^]_ex_ is below 5–10 µM, but reduces the rate at higher [Ca^2+^]_ex_^35, 40, 41^. The research efforts gradually came to a halt in the 1990s, leaving several mechanistic questions unanswered. First, does spermine actually modulate uniporter activity? If yes, what are the underlying molecular mechanisms? Moreover, how do we reconcile the contradictory observations about the kinetics of mitochondrial Ca^2+^ uptake mentioned above?

We decided to pursue these questions for two reasons. First, from the perspective of pure channel biophysics, we are curious about how spermine potentially enhances uniporter function, instead of just blocking the pore as in the case of other cation channels. Second, spermine regulation of mitochondrial Ca^2+^ buffering has been considered as physiologically relevant as the half maximal effective concentration (EC_50_) of spermine, 100–200 µM, falls into the concentration range of free spermine in the cytoplasm^44^. Thus, a deep understanding of spermine potentiation could advance our knowledge of how cells regulate the all-important processes of cytoplasmic and mitochondrial Ca^2+^ signaling and homeostasis.

## Results

### Spermine perturbs MICU1 block of the uniporter

To observe the potentiation of mitochondrial Ca^2+^ buffering by spermine, wild-type (WT) HEK293 cells were permeabilized with digitonin in the presence of Fluo-4 to monitor changes in [Ca^2+^]_ex_. Fig. 2A shows that mitochondria reduce [Ca^2+^]_ex_ to a steady-state level of 517 ± 27 nM, and adding 1 mM of spermine causes a further reduction of [Ca^2+^]_ex_ by about 50% (255 ± 8 nM). Experiments relating spermine concentrations to percentage of [Ca^2+^]_ex_ reduction yielded an EC_50_ of 201 µM (Fig. 2B). These results are consistent with the original data from Nicchitta and Williamson^35^. We also show that following spermine addition, increased mitochondrial Ca^2+^ absorption is indeed mediated by the uniporter, as this process can be abolished by using Ru360 to inhibit the uniporter (black trace, Fig. 2A).

**Figure 2.**
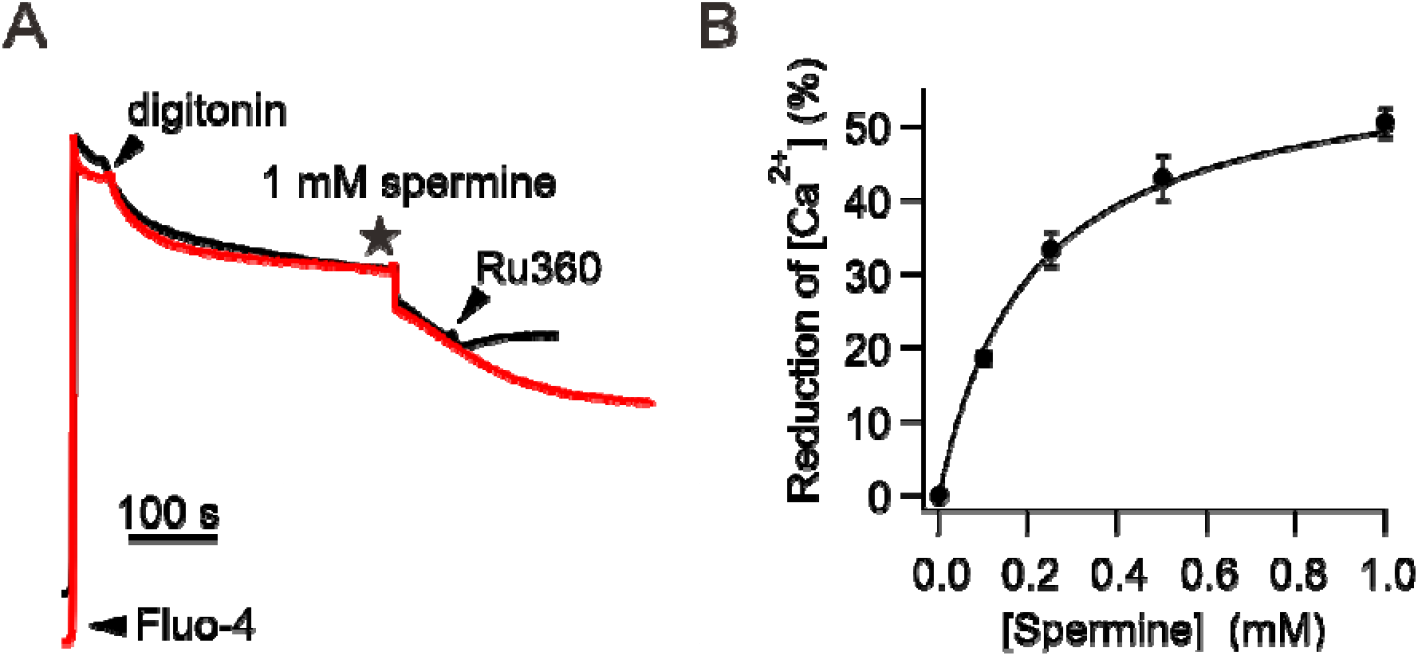
Enhancement of mitochondrial Ca^2+^ absorption induced by spermine. **(A)** Spermine potentiation of mitochondrial Ca^2+^ uptake. Adding 1 mM spermine causes mitochondria to sequester more Ca^2+^. This spermine effect is abolished by adding 100 nM Ru360 as shown in the black trace. **(B)** Spermine dose-response relationship.

The effect of spermine on [Ca^2+^]_ex_ is remarkably similar to that caused by MICU1-knockout (KO) — In the absence of MICU1, mitochondria can absorb more Ca^2+^ to achieve lower [Ca^2+^]_ex_ because the uniporter would not be blocked by MICU1 when [Ca^2+^]_ex_ decreases (Fig. 1A). Such similarity led us to hypothesize that spermine exerts its potentiation effect by perturbing MICU1 block of the MCU pore. This hypothesis predicts that spermine would pose no effects on mitochondria Ca^2+^ absorption when the uniporter is not blocked by MICU1 under the following conditions: (1) the MICU1 gene is deleted (MICU1-KO), (2) WT MICU1 is substituted by a K126A MICU1 mutant that cannot block MCU^12^, or (3) MICU1 is separated from the MCU pore due to increased [Ca^2+^]_ex_. Indeed, we show that MICU1-KO abolishes spermine’s stimulatory effects (Fig. 3A-B), and the phenotype can be restored by expressing WT but not K126A-MICU1 (Fig. 3B-C). Moreover, while spermine increases the rate of uniporter-mediated mitochondrial Ca^2+^ uptake at 1-µM [Ca^2+^]_ex_, the effect becomes smaller at 5-µM [Ca^2+^]_ex_, and is completely eliminated when [Ca^2+^]_ex_ rises to 10 µM (Fig. 3D-E). Altogether, these results demonstrate that spermine enhances mitochondrial Ca^2+^ buffering by inhibiting MICU1 block of the uniporter. Of note, we observed no effects of spermine on inner mitochondrial membrane (IMM) potentials (Fig. 3F), indicating that spermine does not affect the driving force for Ca^2+^ influx, a conclusion in line with previous reports^35, 42^.

**Figure 3.**
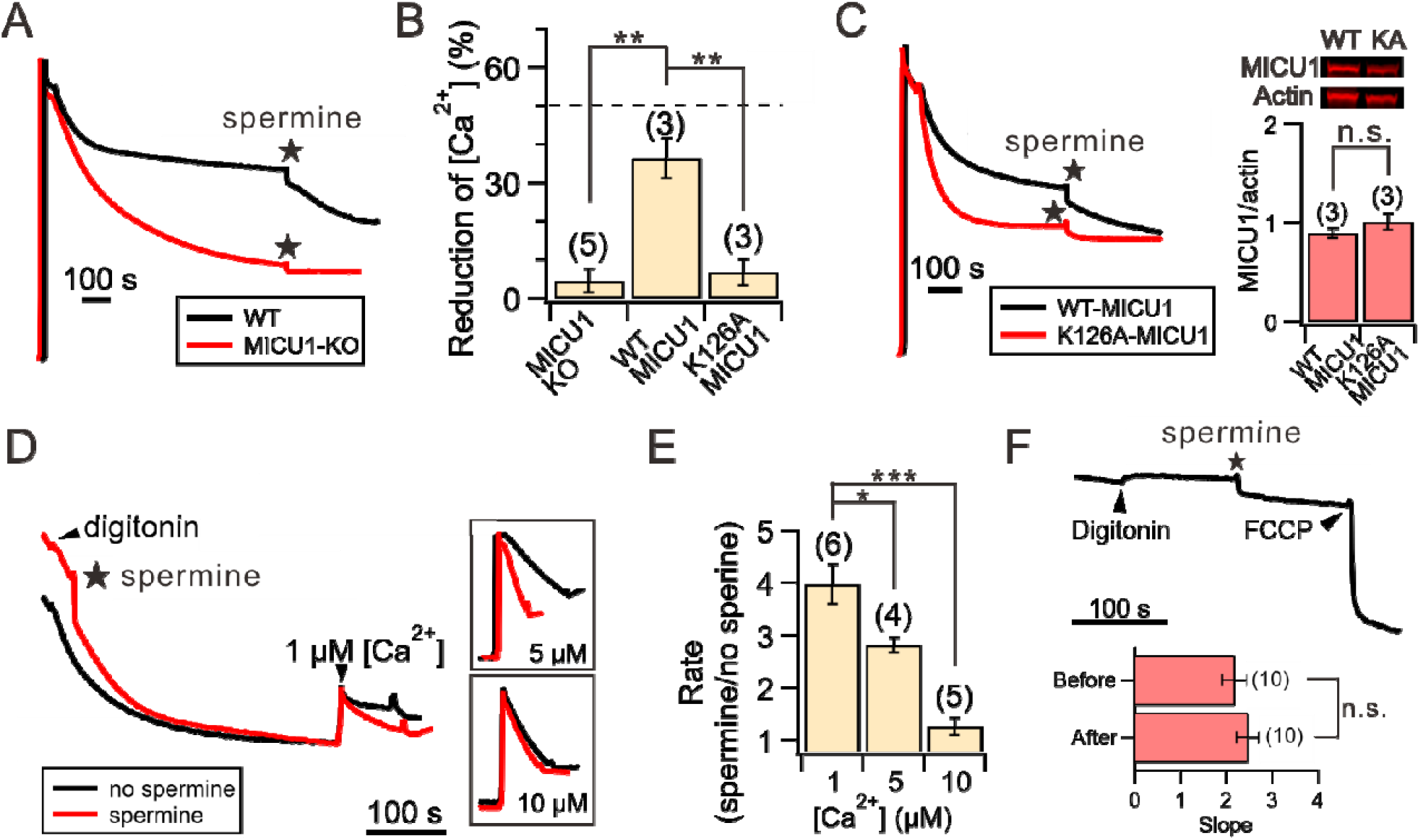
Spermine perturbation of MICU1 block. **(A)** MICU1 dependence of spermine potentiation effects. **(B)** A bar chart summarizing the ability of spermine to induce reduction of [Ca^2+^]_ex_ in MICU1-KO cells (left), or MICU1-KO cells transfected with WT MICU1 (middle) or K126A MICU1 (right). *Dashed line*: WT HEK cells. **(C)** Representative traces of spermine effects on MICU1-KO cells transfected with WT or K126A MICU1. The two MICU1 constructs were expressed to similar levels as shown in Western blot images, which were quantified in the bar chart. **(D)** Spermine-induced acceleration of mitochondrial Ca^2+^ uptake. Various concentrations of Ca^2+^ were added to induce net mitochondrial Ca^2+^ uptake in the presence (red) or absence (black) or spermine. Inlets highlight the traces after adding Ca^2+^. **(E)** Fold-increase of mitochondrial Ca^2+^ uptake rate in response to spermine. **(F)** Lack of spermine effects on IMM potentials. The slopes before and after adding spermine are presented in the bar chart. 1 mM spermine was used in the entire figure. *P < 0.05; **P < 0.01; ***P < 0.001; n.s.: no significance.

### The mechanisms underlying spermine potentiation

The results above indicate that spermine potentiation requires at least MCU, EMRE, and MICU1 because the effect can be eliminated by (1) MICU1-KO (Fig. 3A-B) or (2) using Ru360 to block Ca^2+^ influx through a Ca^2+^-conducting pore formed by MCU and EMRE (Fig. 2A). As in HEK cells, the uniporter also possesses MICU2, which resides in a MICU1-MICU2 heterodimer (MICU1-2) (Fig. 1A), we asked if MICU2 is necessary for spermine’s actions. Accordingly, we applied spermine to permeabilized MICU2-KO HEK cells, whose uniporters contain a MICU1-MICU1 homodimer (MICU1-1)^9, 45^. Mitochondria in these cells reduce [Ca^2+^]_ex_ to 394 ± 43 nM, and 1 mM of spermine causes a further reduction to 211 ± 15 nM (Fig. 4A). The EC_50_ is 158 µM (Fig. 4B), comparable with that obtained using WT cells (Fig. 2B). We thus conclude that MICU2 is not required for spermine potentiation. Previous studies^9, 19^ suggest that MICU1-1 separates from MCU more readily than MICU1-2. It is thus anticipated that spermine disrupts MCU block by MICU1-1 more easily than MICU1-2. This is indeed the case, as exponential fit of the [Ca^2+^]_ex_ reduction time course following spermine addition (blue curves, Fig. 4A) shows that a new steady state is reached ∼2-fold faster in MICU2-KO than in WT cells (Fig. 4C).

**Figure 4.**
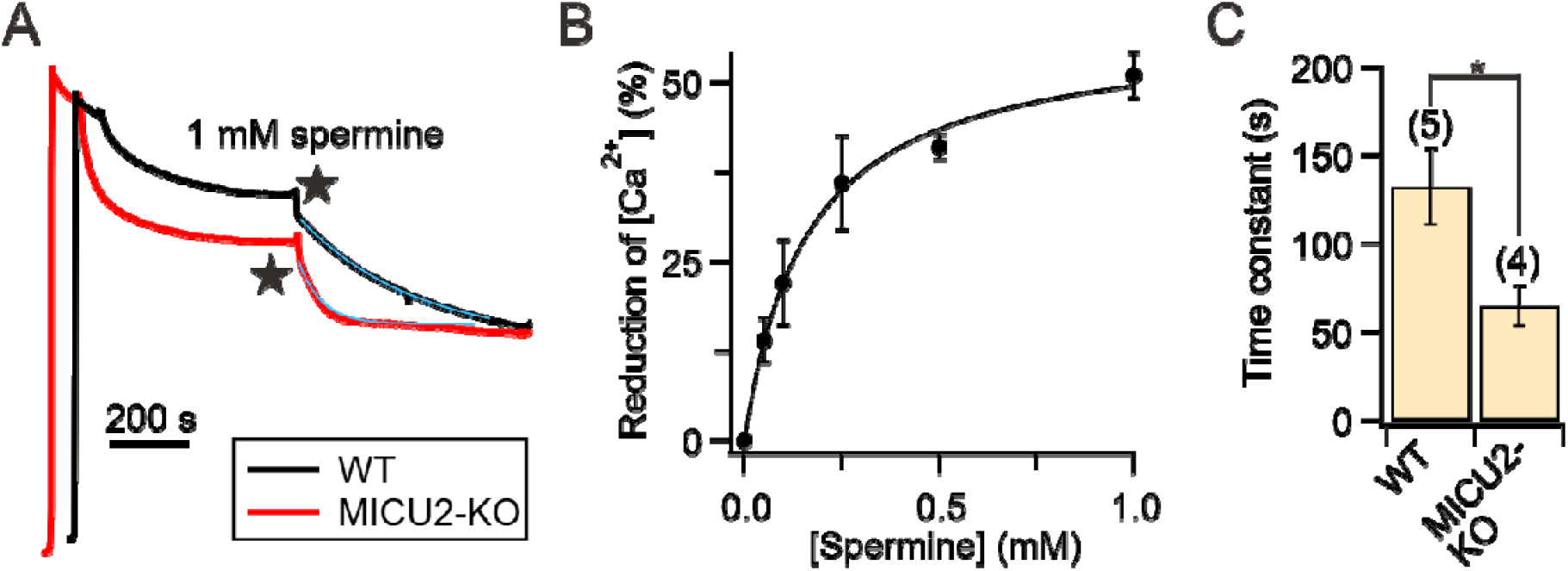
Spermine effects on MICU2-KO cells. **(A)** Representative traces comparing spermine actions on WT vs. MICU2-KO cells. Blue curves: single exponential fit. **(B)** Spermine dose-response relationship obtained using MICU2-KO cells. **(C)** The time constants of spermine-induced [Ca^2+^]_ex_ reduction.

We considered two possible mechanisms about how spermine perturbs MICU1 block. First, spermine prevents MICU1 from binding to MCU to occlude the pore. Second, spermine might disrupt the EMRE-MICU1 interaction, thus facilitating MICU1 dissociation from the uniporter complex^15^. To test these alternative scenarios, we conducted co-immunoprecipitation (CoIP) experiments examining how spermine affects MCU-MICU1 or EMRE-MICU1 interactions. Results show that MICU1 binding to EMRE is unaffected by 1 mM spermine (Fig. 5A). By contrast, spermine fully breaks complex formation between MCU and the MICU1-1 dimer, recapitulating the effect of adding 10 µM Ca^2+^ to dislodge MICU1-1 from MCU (Fig. 5B). We thus conclude that spermine enhances mitochondrial Ca^2+^ absorption by breaking the MCU-MICU1 interactions that block the uniporter.

**Figure 5.**
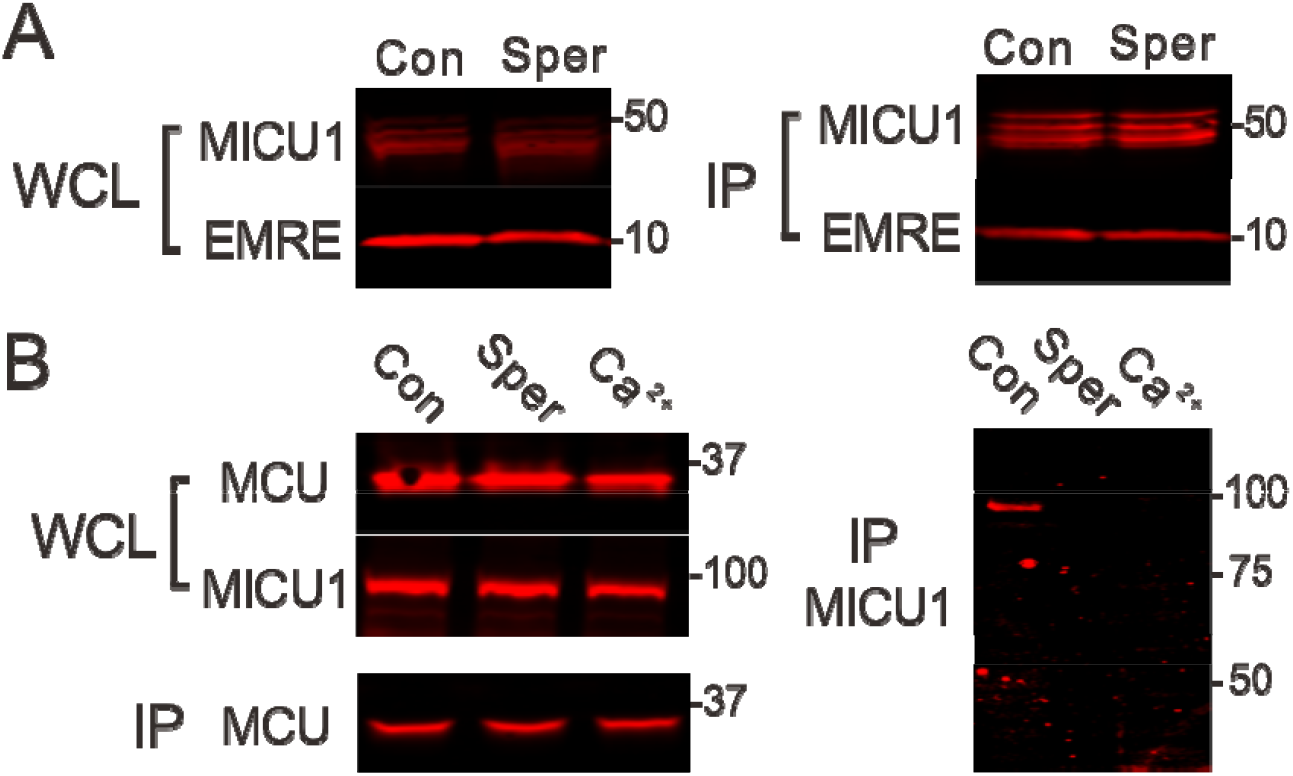
Perturbation of uniporter subunit interactions. **(A)** Spermine effects on MICU1-EMRE interactions. Indicated uniporter subunits were expressed in MICU1-MCU-EMRE-KO cells. MICU1 was used to pull down EMRE. A C463S MICU1 mutant, which cannot form a disulfide-connected MICU1-1 dimer, was used in this experiment, but using WT MICU1 produces similar results that spermine does not affect MICU1 binding to EMRE. **(B)** Disruption of the MCU-MICU1 complex by spermine. MCU was immobilized to pull down WT MICU1 coexpressed in MICU1-MCU-EMRE-KO cells. The MICU1 band migrates to ∼100 kDa as it represents a MICU1-1 dimer connected by an intersubunit disulfide. *WCL*: whole-cell lysate. *IP*: proteins obtained after CoIP. *Con*: 1 mM EGTA. *Sper*: 1 mM EGTA and 1 mM spermine. *Ca*^*2+*^: no EGTA and 10 µM added Ca^2+^. N = 3 for all experiments.

Where does spermine bind? We fist consider an idea that spermine might bind to MICU1’s two EF-hand Ca^2+^ binding motifs to drive MICU1 into a conformation that mimics the Ca^2+^-bound conformation that cannot block MCU. Accordingly, we tested spermine effects on a MICU1 mutant (ΔEF MICU1) containing 4 mutations (D231A, E242K, D421A, and E423K) to disable both EF hands. However, Fig. 6 shows that 1 mM spermine causes a similar reduction of [Ca^2+^]_ex_ in MICU1-KO cells expressing WT or ΔEF MICU1. This result thus ruled out the possibility that spermine binds to EF hands. We are currently in the process of identifying the spermine binding site.

**Figure 6.**
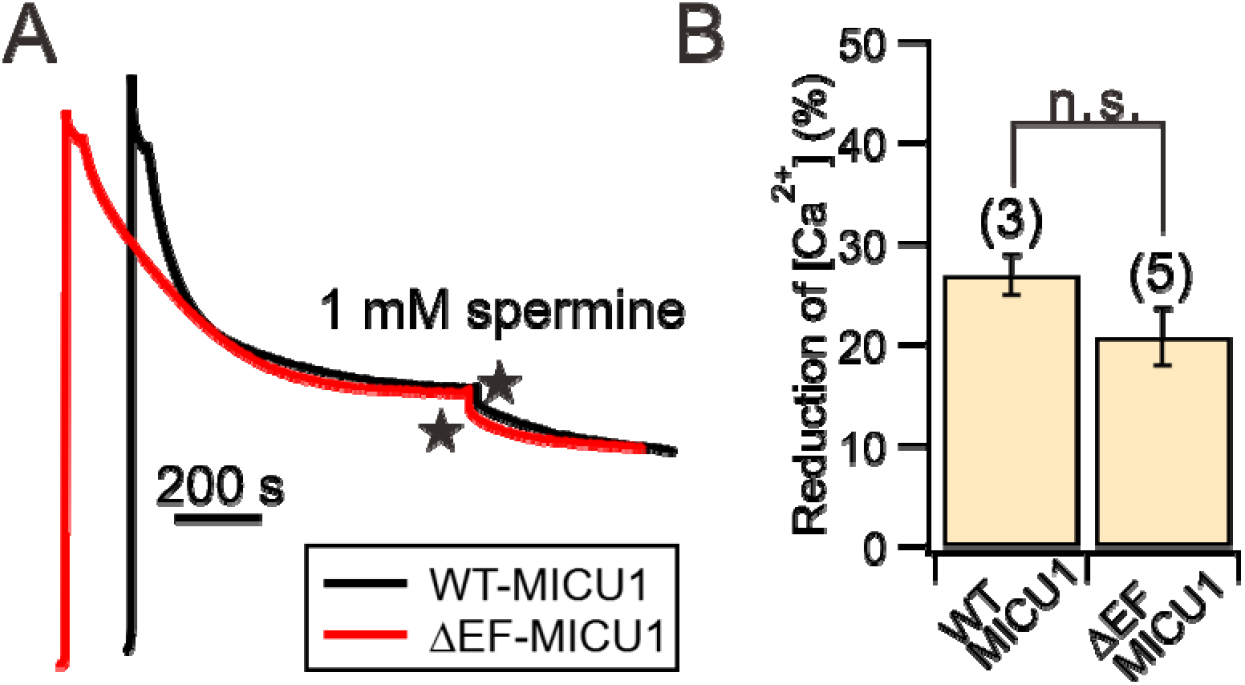
The effect of disabling MICU1 EF hands on spermine action. **(A)** Representative traces of spermine potentiation on MICU1-KO cells transfected with WT- or EF-hand disabled (ΔEF) MICU1. **(B)** A bar chart summarizing the reduction of [Ca^2+^]_ex_ as in (A).

### Spermine also blocks the uniporter

We then sought to examine if spermine also inhibits the uniporter, because spermine is well known for its function of blocking cation channels^24^, and as it has been shown previously that spermine can suppress mitochondrial Ca^2+^ uptake under some conditions^35-37, 40, 41, 43^. To this end, various concentrations of spermine were added to permeabilized WT HEK cells. After [Ca^2+^]_ex_ reaches a steady state, 10 µM Ca^2+^ was then added to fully release MICU1 block. When MICU1 block is relieved, spermine loses its potentiation effects (as shown in Fig. 3D-E). Therefore, the initial rate of mitochondrial Ca^2+^ uptake immediately after adding 10 µM Ca^2+^ would inform us whether spermine can inhibit the uniporter. Fig. 7A shows that with 1-mM spermine, no inhibition was detectable. However, uniporter Ca^2+^ uptake is reduced by ∼30% and ∼45% when [spermine] is increased to 2.5 and 7.5 mM, respectively (Fig. 7A). These results thus demonstrate that spermine also exerts inhibitory effects on the uniporter, albeit with a >10-fold lower potency than the stimulatory effect.

**Figure 7.**
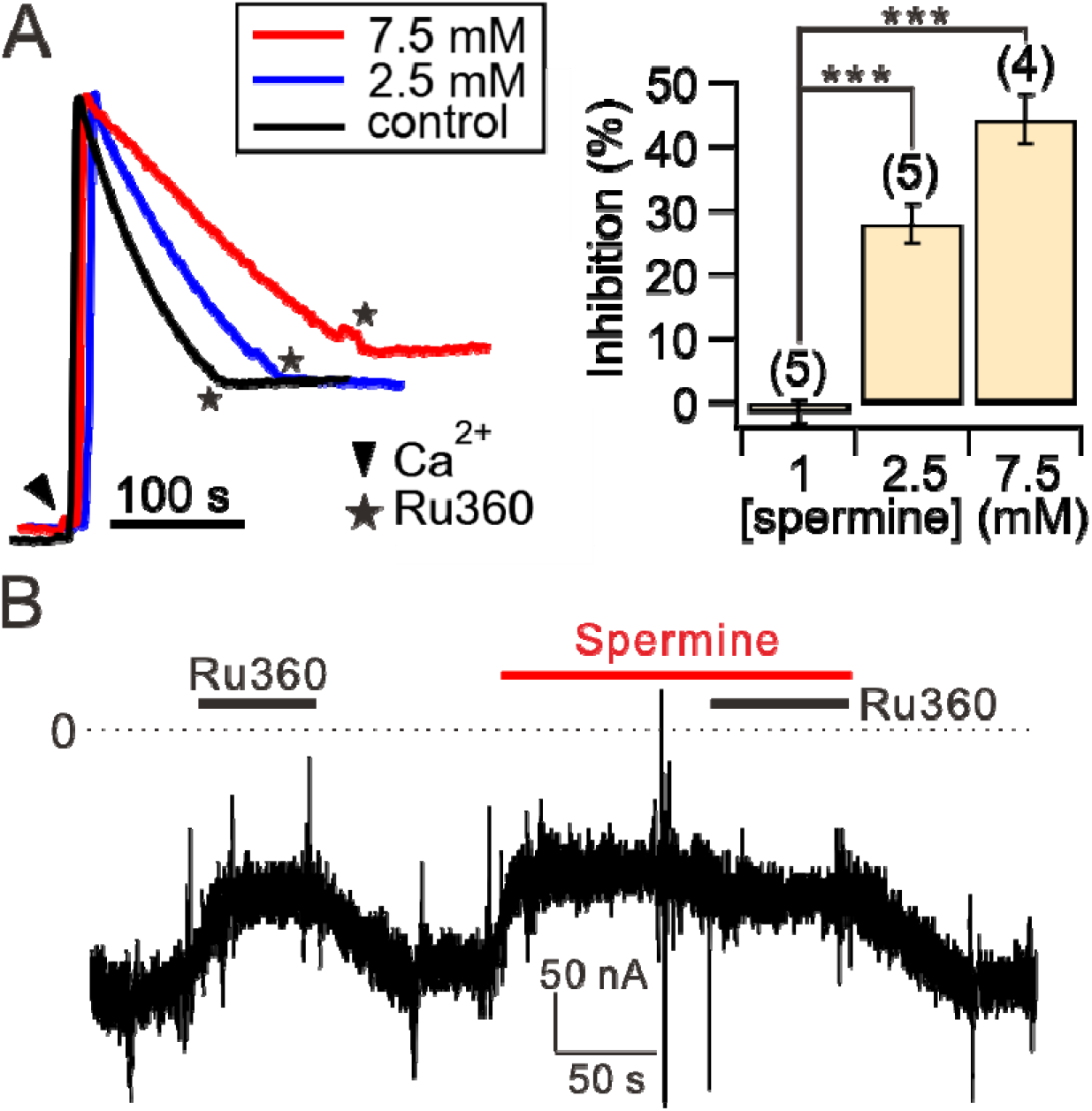
Spermine inhibition of the uniporter. **(A)** Spermine inhibition of mitochondrial Ca^2+^ uptake. The initial rate of mitochondrial Ca^2+^ uptake in the presence of 1, 2.5, or 7.5 mM spermine was normalized to that in the absence of spermine (control) to calculate the percentage of inhibition as shown in the bar chart. **(B)** A representative TEVC recording showing spermine inhibition. 100 nM of Ru360 was added first to identify hME-mediated inward Ca^2+^ current. Applying spermine causes a similar level of current reduction as induced by Ru360, suggesting that spermine fully inhibits the uniporter. Consistently, Ru360 does not further suppress the current when applied in the presence of spermine. *Dashed line*: 0 current.

To further investigate the mechanisms underlying spermine inhibition, we used two-electrode voltage clamp (TEVC) to assess spermine effects on a human MCU-EMRE fusion protein (hME) expressed in *Xenopus* oocytes^46^. We recorded hME-mediated inward Ca^2+^ currents with voltage clamped at -80 mV in the presence of 20 mM Ca^2+^ in the perfusion solution. As expected, varying [spermine] from 50 µM to 5 mM produces no potentiation effect (not shown) since no MICU1 is present in the system. By contrast, hME is fully inhibited by low millimolar levels of spermine (Fig. 7B). These results demonstrate that spermine inhibits the uniporter by targeting the transmembrane MCU-EMRE region. Ongoing electrophysiological work assessing voltage dependence of the inhibition could help address the issues of whether spermine acts as a pore blocker and why spermine inhibition appears to be more potent in oocytes than HEK cells.

## Discussion

This work demonstrates that spermine potentiates the uniporter by perturbing MICU1 block of MCU, while inhibits the uniporter by targeting MCU-EMRE in an MICU-independent manner. Indeed, observations made decades ago have already hinted about the mechanisms of spermine potentiation. It is well known that as [Ca^2+^] increases, the rate of mitochondrial Ca^2+^ uptake rises along a sigmoidal curve, as increased [Ca^2+^] relieves MICU1 block of MCU in a cooperative manner to open the uniporter. Kröner reported that spermine, similar to MICU1-KO^9, 16, 47^, altered this Ca^2+^ dose-response from a sigmoidal to a linear relationship^40^, consistent with spermine abolishing MICU1’s regulatory effect as proposed here.

Our results could help explain some puzzling observations in the literature. It was reported that spermine stimulates mitochondrial Ca^2+^ uptake in hepatic mitochondria but not cardiac mitochondria^40^ (but also see another reference^42^). This observation is consistent with our recent findings that cardiac mitochondria have very low levels of MICU1, leading to a large population of MICU1-free uniporters^9^. These MICU1-deregulated uniporters exhibit a linear [Ca^2+^] dose-response as we and others observed^9, 48^, and are expected to show no response to spermine. As described in the Introduction, previous studies reported complicated effects of spermine on mitochondrial Ca^2+^ uptake kinetics. These can be understood in light of spermine’s dual, independent modulatory effects on the uniporter. As the inhibitory effect requires higher [spermine], it explains why multiple groups^35, 40, 41^ observed spermine potentiation at low [spermine] while the effects become inhibitory as [spermine] rises. Some labs observed only potentiation^39^ or inhibition^36, 37, 43^. This could be due to the concentration of spermine used in their experiments falling into the range in which the stimulatory or inhibitory effects dominate.

An important issue that needs to be addressed regards where spermine binds to modulate uniporter activity. We showed that spermine does not contact the EF hands in MICU1. This is not surprising as EF hands are commonly present in other Ca^2+^ binding proteins, and there have been no reports to date that spermine can interact with these EF hands. This work shows that spermine can inhibit the uniporter in the absence of MICU1 dimers. We speculate that spermine acts as a pore blocker, similar to how it inhibits other cation ion channels. We are conducting further studies to complete our knowledge about spermine regulation of the uniporter complex.

## Materials and methods

### Cell Culture and Molecular Biology

HEK293 cells were cultured in Dulbecco’s Modified Eagle Medium (DMEM) with 10% FBS in a 5% CO_2_ incubator at 37 °C. Deletion of uniporter-subunit genes in HEK cells was achieved via CRISPR-Cas9 as described in our previous work^15, 49^. Site-directed mutagenesis was performed using the QuickChange kit (Agilent), with sequences verified using Sanger sequencing.

### CoIP and western blots

To perform CoIP, HEK cells in 10-cm dishes were transfected at 70-80% confluency and used for experiments 2 days after transfection. Cells were lysed in the presence of a protease inhibitor cocktail (Roche cOmplete EDTA-free) in 1 mL of an ice-cold solubilization buffer (SB, 150 mM NaCl, 50 mM HEPES, 4 mM DDM, pH 7.4-NaOH) supplemented with 1 mM EGTA, 1 mM spermine, or 10 μM CaCl_2_. 10 minutes after cell lysis, the lysate was clarified by spinning down at 13,000 g for 10 min. 100 μL of the lysate was taken out and used as the whole-cell lysate samples in Fig. 5. 25 μL of FLAG-conjugated (Sigma Aldrich, A2220) or 1D4-conjugated (homemade, 50% slurry) resins were then added to the remaining cell lysate and incubated on a tube revolver at 4°C for 1 hour. The beads were collected on a spin column, washed six times with 1-mL SB, and then eluted with 140 μL of 1X SDS loading buffer for the IP samples in Fig. 5.

Protein samples were transferred to low-fluorescence PVDF membranes and were blocked in a Tris-buffered saline (TBS)-based intercept blocking buffer (Li-Cor). The membranes were incubated with primary antibodies diluted in TBST (TBS plus 0.075% Tween-20) at 4□°C overnight. These include: anti-1D4 (produced in house, 0.1 mg/mL), anti-MICU1 (Sigma Aldrich HPA037480, 1:10,000), or anti-FLAG (Sigma Aldrich F1804, 1:10,000) antibodies. After 1-hour incubation with fluorescent secondary antibodies, goat anti-rabbit IRDye 680RD (Li-Cor 92568171, 1:10,000) or goat anti-mouse IRDye 680RD (Li-Cor 925-68070, 1:15,000), at room temperature, the signals were captured using a LI-COR Odyssey CLx imager. Band intensities were quantified using the LI-COR Image Studio software (version 5.2).

### Mitochondrial Ca^2+^ uptake assays

To quantify the stimulation effects of spermine, 2 × 10^7^ HEK cells were washed in 10 mL of a wash buffer (120 mM KCl, 25 mM HEPES, 2 mM KH_2_PO_4_, 1 mM MgCl_2_, 50 mM EGTA, pH 7.2-KOH), and resuspended in 2 mL of a recording buffer (120 mM KCl, 25 mM HEPES, 2 mM KH_2_PO_4_, 1 mM MgCl_2_, 5 mM succinate, pH 7.2-KOH). The sample was placed in a stirred quartz cuvette in a Hitachi F-7100 spectrofluorometer (ex: 494nm, ex-slit: 2.5 nm, em: 516 nm, em-slit: 5.0 nm, sampling frequency: 1 Hz). After adding 0.25 μM of Fluo-4 (Invitrogen, F14200) to monitor [Ca^2+^]_ex_, 30 mM of digitonin (Sigma, D141) was added to permeabilize the cells. When [Ca^2+^]_ex_ reached a steady state, spermine was added to induce further mitochondrial Ca^2+^ absorption. At the end of the recording, 40 μM of CaCl_2_ was added to obtain the saturating fluorescence (F_max_), followed by adding 500 μM of EGTA to obtain the minimum fluorescence signal (F_min_). [Ca^2+^]_ex_ at a certain fluorescence signal (F) was calculated using the following equation using a Fluo-4 Ca^2+^ K_d_ of 345 nM as provided by the manufacturer:

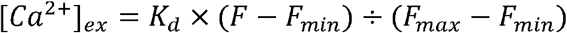

To measure the rate of mitochondrial calcium uptake rate (Fig. 3D-E and 7A), Calcium Green 5N (Invitrogen, C3737) was used to monitor [Ca^2+^]_ex_. Quantification was done by calculating the slope of the first 10 s of fluorescence signal decline after adding Ca^2+^.

### IMM depolarization essay

2 × 10^7^ HEK cells were incubated with 40 nM of TMRM (Invitrogen, T668) in Tyrode’s solution (130 mM NaCl, 5.4 mM KCl, 1 mM MgCl_2_, 1 mM CaCl_2_, 20 mM HEPES, pH 7.8-NaOH) at 37 °C for 30 min. Cells were washed and pelleted down using 10 mL of a Mg^2+^-free wash buffer (120 mM KCl, 2 mM K_2_HPO_4_, 50 μM EGTA, 25 mM HEPES, pH 7.2-KOH) and resuspended using a 2 mL Mg^2+^-free recording buffer (120 mM KCl, 2 mM K_2_HPO_4_, 5 mM succinate, 25 mM HEPES, pH 7.6-KOH). The sample was loaded in a stirred quartz cuvette in a Hitachi F-7100 spectrofluorometer (excitation: 573 nm; excitation slit: 5 nm; emission: 590 nm; emission slit: 5 nm; sampling rate: 1 Hz). After the cells were permeabilized by 30 mM of digitonin, 1 mM of spermine was added to the cuvette, followed by adding 1 μg/mL of FCCP to completely collapse the IMM potential. The rate of IMM depolarization was quantified by normalizing the slope before or after adding spermine to the total TMRM signal.

### Electrophysiology

TEVC was performed as described before^46^. Briefly, stage V-VI oocytes were injected with 40 ng of hME cRNA and incubated in 18°C in a ND96 solution (96 mM NaCl, 2 mM KCl, 2 mM CaCl_2_, 0.5 mM MgCl_2_, 5 mM HEPES, pH 7.4-NaOH). Recordings were performed 2–3 days after cRNA injections. Signals were measured using the Oocyte Clamp OC-725B system (Warner). Voltage and current electrodes were filled with 3 M KCl to have a resistance of 0.5–1 MΩ. All recordings were performed in a Ca-20 solution (70 mM NaCl, 2 mM KCl, 0.5 mM MgCl_2_, 20 mM CaCl_2_, 5 mM HEPES, pH 7.4-NaOH).

### Statistics

Statistical analysis was performed using two-tailed t test with significance defined as P < 0.05. We performed at least three independent repeats in all experiments. Data are presented as means ± S.E.M. Electrophysiology and mitochondrial Ca^2+^ flux data were analyzed using Igor Pro 8 (WaveMetrics). All figures were prepared using CorelDRAW.

## Acknowledgements

This work is supported by the NIH grant R01-GM129345.

## Author Contributions

MFT conceptualized the project. YCT and FYC performed experiments and analyzed data. MFT and YCT prepared the manuscript.

